# A web-based, branching logic questionnaire for the automated classification of migraine

**DOI:** 10.1101/369827

**Authors:** Eric A. Kaiser, Aleksandra Igdalova, Geoffrey K. Aguirre, Brett Cucchiara

## Abstract

**Objective:** To identify migraineurs and headache-free individuals with an online questionnaire and automated analysis algorithm.

**Methods:** We created a branching-logic, web-based questionnaire—the Penn Online Evaluation of Migraine (POEM)—to obtain standardized headache history from a previously studied cohort. Responses were analyzed with an automated algorithm to assign subjects to one of several categories based on ICHD-3 (beta) criteria. Following a pre-registered protocol, this result was compared to prior diagnostic classification by a neurologist following a direct interview.

**Results:** Of 118 subjects contacted, 90 (76%) completed the questionnaire; of these 31 were headache-free, 29 migraine without aura (MwoA), and 30 migraine with aura (MwA). Mean age was 41 ± 6 years and 76% were female. There were no significant demographic differences between groups. The median time to complete the questionnaire was 2.5 minutes. Sensitivity of the POEM tool was 42%, 59%, and 70%, and specificity was 100%, 84%, and 94% for headache-free, MwoA, and MwA, respectively. Sensitivity and specificity of the POEM tool for migraine overall (with or without aura), was 83% and 90%, respectively.

**Conclusions:** The POEM web-based questionnaire, and associated analysis routines, identifies headache-free and migraine subjects with good specificity. It may be useful for classifying subjects for large-scale research studies.

**Trial Registration:** https://osf.io/sq9ef

## Introduction

Migraine is a highly prevalent, complex neurologic disorder (1), and includes variants with and without antecedent aura. Research studies of migraine require a means to classify subjects by headache type and to identify headache-free subjects as a comparison group. Traditionally, a clinical interview by a neurologist is used for this purpose, often guided by the diagnostic criteria of the International Classification of Headache Disorders (ICHD)(2). While considered the “gold standard”, a limitation of this approach is that it can be time-consuming, resource-limited, and difficult to standardize.

Numerous screening tools have been proposed as an alternative means to identify patients with migraine. Many of these are designed to screen for migraine within the primary care setting (3–11), and involve 3 to 5 questions regarding duration of, or disability from, headaches and associated headache features including nausea, photophobia, and/or phonophobia. Several surveys have been designed specifically for research purposes (1, 12, 13). While these tools can identify migraineurs, they generally are not designed to distinguish between migraine with aura (MwA) and migraine without aura (MwoA) (except for DMQ-3, which is no longer available online) (13), do not identify headache-free individuals, and are manually administered and analyzed.

Here we present a web-based questionnaire and automated analysis code that applies the ICHD-3 (beta) criteria to subject responses; we dub the instrument the “Penn On-line Evaluation of Migraine”, or POEM. The goal is to provide an efficient means to classify subjects as migraineurs or headache-free controls. The questionnaire makes use of branching logic to minimize test duration and the presentation of irrelevant questions. The resulting questionnaire data are processed with unit-tested, open-source routines that automate headache classification. To validate the POEM, we identified subjects who had previously been assigned a headache classification (including the classification of headache-free) by one of us as part of a prior research study (14). Following a pre-registered protocol (https://osf.io/sq9ef), we contacted these subjects and invited them to complete the POEM. We then compared the headache classification provided by the POEM with their prior classification, and computed the sensitivity and specificity of the test.

## Methods

### Ethics statement

The study was approved by the University of Pennsylvania Institutional Review Board; all participants provided written informed consent for participation in the original study from which subjects were drawn. For assessment of the web-based questionnaire, a waiver of written consent was granted by our Institutional Review Board.

### Design of the POEM instrument

Using a commercially available survey tool (http://qualtrics.com/), we created a branching-logic, web-based questionnaire that guides subjects through a series of questions regarding headache history and associated symptoms. The question text and available responses were designed to assign patients to the ICHD-3 (beta) categories of MwoA or MwA; and to further distinguish between visual and non-visual aura. In addition, questions attempted to differentiate between individuals who are headache-free or experience headaches that are mild and have non-migrainous features (mild non-migrainous headache) or are more severe and/or have at least one migrainous feature (headache not otherwise specified, NOS). Subjects who endorsed photophobia were also administered a set of questions regarding their ictal photophobia symptoms taken from a prior report (15). Finally, the survey included questions regarding headache recency, medication use, childhood motion sickness, menstrual headaches, and family history of migraine. This final set of questions did not inform the assignment of diagnostic category described here.

As a validity check, subjects were asked “Did you take this survey seriously and are your answers sincere? Are all your answers correct?”, and offered three responses 1) “Yes, I took the survey seriously and my answers are correct”; 2) “I made an error and want to retake the survey”; “I didn’t take this seriously. Don’t use my answers”. If a subject responded with the second response, they were invited to retake the survey, and the second set of responses were retained. If they failed to successfully retake the survey, they were excluded from the study. No subjects replied with the third response. For an illustration of the branching-logic scheme, see Figure 1; for a complete list of the survey questions and available responses, see Supplemental Figure 1.

**Figure 1.**
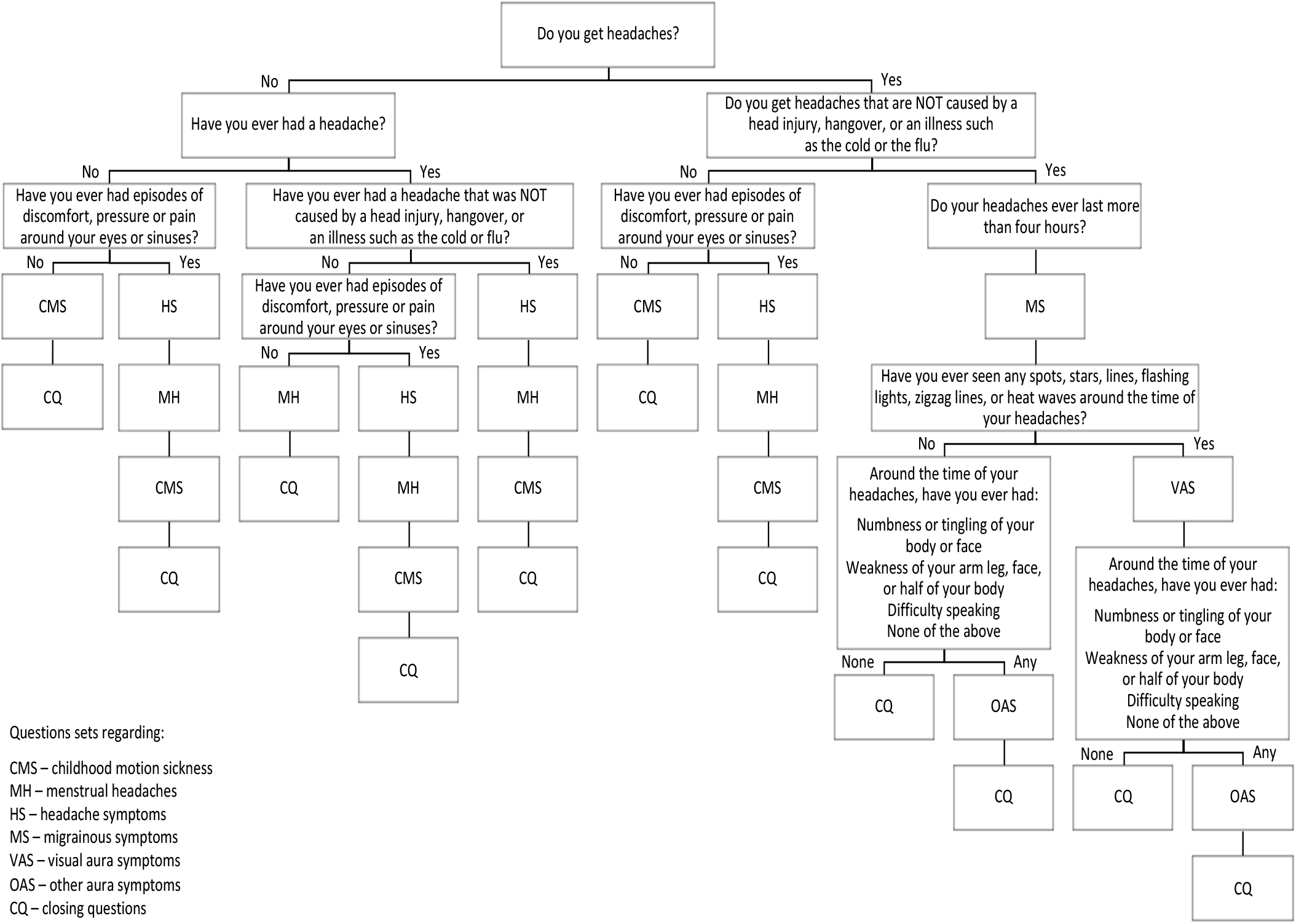
Diagram of POEM questionnaire

Categorical assignment was performed using automated analysis routines implemented in MATLAB (https://github.com/gkaguirrelab/poemAnalysis). While the code (and associated unit tests) provide the canonical description of the algorithm, we outline here the principles followed for categorical assignment. Inclusion criteria for MwoA, migraine with visual aura, and migraine with other aura are outlined in Supplemental Tables 1-3 and are based on ICHD-3 (beta) criteria. Note that subjects could potentially receive multiple migraine classifications. For headache-free, subjects can meet criteria based on three potential pathways (A, B, or C) within the branching logic structure (Supplemental Table 4) in which they deny any headaches or facial pain/discomfort/pressure or deny having headaches that were not be caused by head injury, hangover, or illness. If subjects did not meet criteria for headache-free, subjects could be considered for mild non-migrainous headache unless their responses meet any exclusion criteria that suggest moderate to severe headaches and/or headaches with at least one migrainous feature (Supplemental Table 5). Finally, if subjects do not meet criteria for any of the prior classifications, subjects are given a classification of headache NOS.

### Validation study

Subjects were recruited from a prior observational cohort, the Anatomy and Cerebral Hemodynamic Evaluation of Migraine (ACHE-M) study, which included both migraine and control subjects. A detailed description of ACHE-M has been published previously (14); briefly, this was a prospective case-control study using magnetic resonance imaging (MRI) to compare vascular structure and function between headache-free controls and MwA and MwoA patients. Participants were 25–50 years old at the time of ACHE-M enrollment, which occurred between March 2008 and June 2012. Participants were screened and examined by a single board-certified neurologist with expertise in headache to determine headache status; subjects were classified as MwA or MwoA using International Classification of Headache Disorders criteria (edition 2)(16), Subjects classified as headache-free healthy controls could not have any history of potentially migrainous headaches (including ‘sinus’ headaches or any type of headache requiring medication) and could not have tension-type headache unless it occurred exceedingly rarely (defined as less than once a year). Isolated, infrequent provoked headaches associated with excessive alcohol consumption or transient illness other than reported sinusitis was allowed. Family history of migraine was not used to exclude headache-free people.

Subjects from ACHE-M were contacted by email and invited to participate in the current research project; each email include an individualized web link for the on-line questionnaire. Subjects were initially contacted on September 15, 2017. Subjects who failed to respond to the initial email were re-contacted 2 additional times roughly 2 weeks and 4 weeks after the initial email. Following the pre-registered protocol, subjects who failed to complete the entire survey by November 1, 2017 were excluded.

### Data Analysis

Groups of participants were compared using the chi-squared or Fisher’s exact test for dichotomous or categorical variables and the *t*-test or Wilcoxon ranked-sum tests for continuous variables as appropriate. Means and standard deviations or medians and interquartile ranges (IQR) are presented as appropriate. An association was considered significant if *p* < 0.05. All tests were two sided. Statistical analyses were performed using JMP (Version 9, SAS Institute Inc., Cary, NC). Specificity and sensitivity were calculated for each ACHE-M classification group (i.e. MwoA, MWA, control). For such analysis, we collapsed the POEM labels of migraine with visual aura and migraine with non-visual aura into a single category; in addition, the headache free classification included patients with mild, infrequent non-migrainous headache as was used in ACHE-M.

### Data and code availability

The analysis code is open source and available for modification, distribution, and use with attribution (https://github.com/gkaguirrelab/poemAnalysis). The repository includes the questionnaire itself in “qsf” format, along with instructions on how to download and implement the instrument in Qualtrics. The anonymized raw data and code that produced the results presented in the current report are also available (https://github.com/gkaguirrelab/Kaiser_2018_TBD).

### Pre-registration

The study and methodology for analysis were pre-registered prior to data collection and analysis (https://osf.io/sq9ef). There were no deviations from the study protocol.

## Results

### Study subjects

Of the original 172 ACHE-M subjects, valid email addresses were available for 118 subjects, and of these, 90 (76%) completed the POEM questionnaire. Subjects who completed the POEM questionnaire were approximately evenly distributed among the three ACHE-M classification groups. Subject recruitment is illustrated in Figure 2. Clinical characteristics of subjects are described in Table 1. Aside from age, which reflects current age at time of completion of the POEM questionnaire, other subject characteristics were collected at time of enrollment in ACHE-M. There was no significant difference between the control and migraine groups in terms of age, sex, migraine duration since age of onset, blood pressure, or history of smoking. There was no significant difference in use of migraine-prophylactic medication between MwA and MwoA groups.

**Figure 2.**
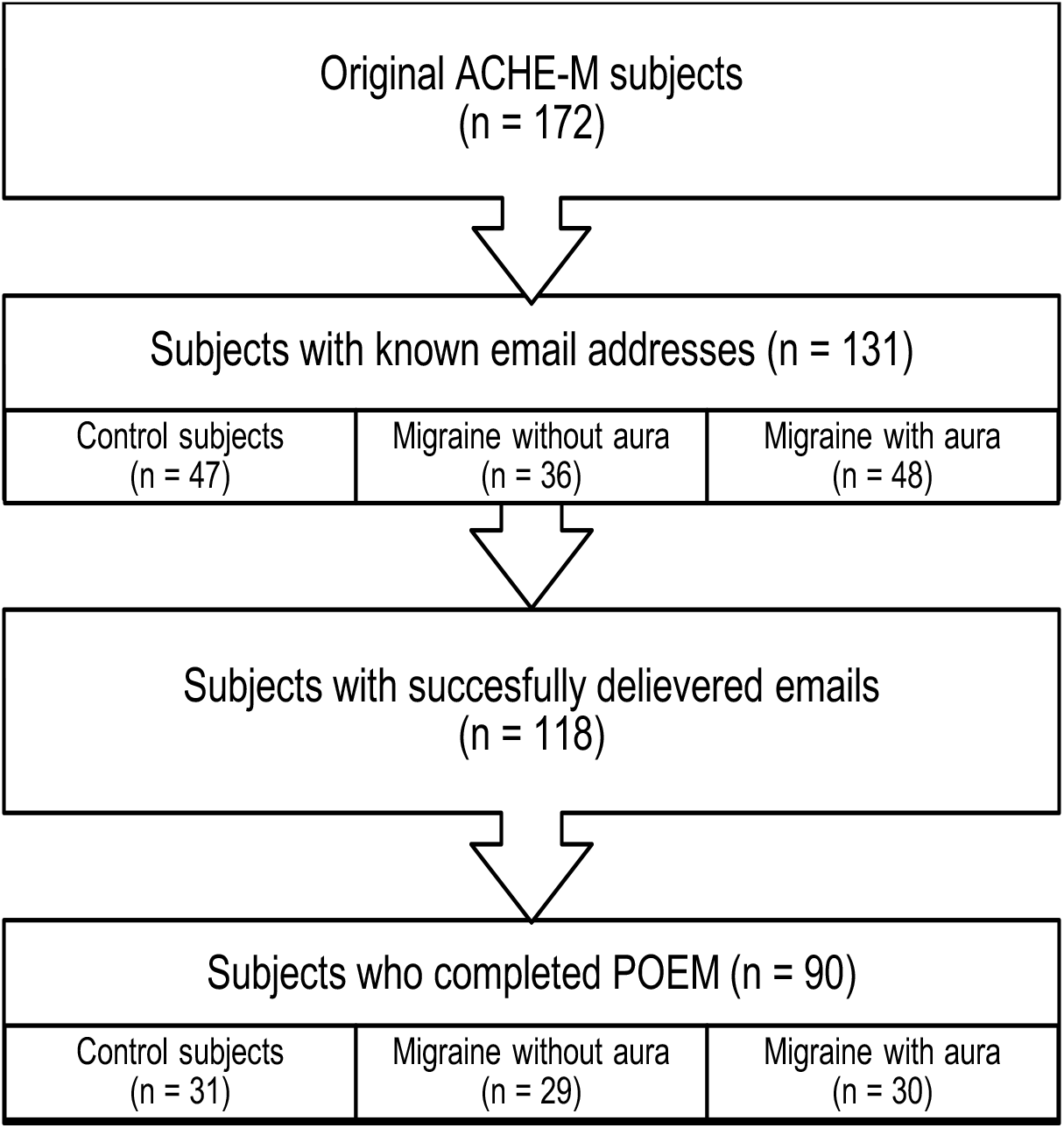
Flow diagram of subject recruitment.

**Table 1.** Clinical characteristics

|  | Control | Migraine<br>wo aura | Migraine w<br>aura | p value |  |  |
| --- | --- | --- | --- | --- | --- | --- |
|  |  |  |  | Control v<br>MwoA | Control v<br>MWA | MwoA v<br>MWA |
| **n** | 31 | 29 | 30 | - | - | - |
| **Age (mean +/- SD)** | 41 ± 7 | 42 ± 6 | 41 ± 6 | 0.75 | 0.81 | 0.57 |
| **Female sex (n, %)** | 21 (68%) | 21 (72%) | 26 (87%) | 0.69 | 0.08 | 0.17 |
| **Migraine duration in years (mean ± SD)** | - | 17 ± 9 | 19 ± 10 | - | - | 0.34 |
| **Migraine frequency, #/month (median, IQR)** | - | 2 (1-5) | 3 (2-6) | - | - | 0.48 |
| **Migraine-prophylactic medication (n, %)** | - | 5 (17%) | 7 (23%) | - | - | 0.56 |
| **Systolic blood pressure (mean ± SD)** | 126 ± 18 | 126 ± 16 | 126 ± 21 | 0.97 | 0.97 | 0.99 |
| **Diastolic blood pressure (mean ± SD)** | 81 ± 10 | 85 ± 12 | 82 ± 14 | 0.15 | 0.65 | 0.42 |
| **Smoker, current or former, n (%)** | 8 (26%) | 11 (38%) | 7 (23%) | 0.31 | 0.82 | 0.22 |

\*Data recorded at time of enrollment in ACHE-M with the exception of age, which was calculated at time of POEM completion based on date of birth.

Completion of the POEM questionnaire required a median time of 2 minutes 32 seconds. Only 1 subject reported making an error during completion of the questionnaire; this subject successfully completed the questionnaire on a second attempt. No subjects indicated they did not take the survey seriously.

### Validation of POEM instrument

A confusion matrix describing diagnostic category in ACHE-M compared to the POEM classification is shown in Table 2. Sensitivity of the POEM instrument was 42%, 59%, and 70% for headache-free, MwoA, and MwA, respectively, and specificity was 100%, 84%, and 94% for headache-free, MwoA, and MwA, respectively (Table 2). Combining MwoA and MwA into a single migraine group, sensitivity and specificity of the POEM instrument was 83% and 90% respectively.

**Table 2.** Confusion matrix comparing classification by direct neurologist interview versus POEM.

|  |  | POEM |  |  |  |  | Total | Sensitivity | Specificity |
| --- | --- | --- | --- | --- | --- | --- | --- | --- | --- |
|  |  | Headache free | Mild non-migrainous headache | Headache NOS | MwoA | MWA |  |  |  |
| ACHE-M | Control | 5 | 8 | 15 | 2 | 1 | 31 | 42% | 100% |
|  | MwoA | 0 | 0 | 9 | 17 | 3 | 29 | 59% | 84% |
|  | MwA | 0 | 0 | 1 | 8 | 21 | 30 | 70% | 94% |
|  | Total | 5 | 8 | 25 | 27 | 25 | 90 |  |  |

We examined the 14 cases in which there was disagreement between the POEM and the original headache classification by the ACHE-M study neurologist. Comparing the misclassified subjects to those correctly classified, there was no significant difference in age, sex, migraine frequency, or years with migraine. Three control ACHE-M subjects now reported symptoms consistent with migraine and were classified by the POEM instrument as having MwA (1 subject) or MwoA (2 subjects). Also, three MwoA ACHE-M subjects now reported aura symptoms and were classified as MwA by the POEM: two endorsed visual aura and one endorsed numbness or tingling. Eight subjects classified as MwA by ACHE-M were classified by the POEM instrument as MwoA. All 8 subjects endorsed visual aura symptoms in the questionnaire but failed to meet strict POEM criteria for MwA for the following reasons: 2 subjects indicated they had aura only once in their lifetime (and thus did not meet criteria for a minimum of 2 lifetime episodes); 1 subject reported having visual aura typically greater than 60 min in duration, and 5 subjects reported having visual aura typically less than 5 min in duration (and thus did not meet criteria for aura symptoms lasting between 5 and 60 minutes).

## Discussion

The POEM is an open-source, automated implementation of the ICHD-3 (beta) criteria for migraine classification. Based on responses from a self-administered, online questionnaire using branching-logic, the POEM instrument classifies subjects into one of six categories: headache-free, mild non-migrainous headache, headache not otherwise specified, migraine without aura, migraine with visual aura, and migraine with other aura. Subjects are generally able to complete the POEM in less than 3 minutes and with rare self-reported errors. The POEM is moderately sensitive but is highly specific in identifying subjects as controls (headache free or mild non-migrainous headache), MwoA, or MwA. It was designed to strictly apply the ICHD-3 (beta) criteria to potential subjects.

We conducted a validation of the POEM by testing a group of subjects who had received a prior headache classification. A significant limitation of this validation effort is the time interval (5-9 years) between classification by the study neurologist for ACHE-M and completion of the POEM questionnaire. This may explain, at least partially, why the sensitivity obtained for the POEM is lower than prior screening tools. While the POEM questions were designed to capture headache and associated symptoms throughout a subject’s lifetime, there is evidence that headache status can change (17–21), and it is likely that recall of prior headache symptoms may degrade over time. Prior epidemiological studies have reported variable reproducibility in the diagnosis of migraine. In one study, migraine diagnosis was reproducible in 78% of the population over 12 months when using strict ICHD criteria (21), but in another study that decreased to only 48% over 2.2 years (17). Another population-based study found that in a 3-year period the incidence of migraine in individuals under 45 years of age increased by a third in women and doubled in men (19). We did not ask subjects to state if or how their headaches may have changed since enrollment in ACHE-M, so we cannot directly address the question of how this contributed to misclassification. However, it is perhaps not surprising that a subset of patients initially classified in ACHE-M as controls went on to develop migraine over the ensuing years.

Of the 14 subjects who were misclassified by the POEM compared to the neurologist’s classification for ACHE-M, 8 of the subjects were initially classified as MwA, but the POEM classified them as MwoA. In all 8 cases, the subjects endorsed aura symptoms but failed to meet ICHD criteria for MwA, although would have met criteria for probable MwA. In 6 cases, the POEM instrument excluded those subjects based on the duration of their symptoms. The specific question posed to subjects was “How long do these phenomena typically last?”. The ICHD criteria do not require that aura phenomena “typically” last between 5 and 60 minutes, only that some episodes do so. We suspect that the particular wording of this question was responsible for the overly conservative behavior of the POEM for the diagnosis of MwA. Using a more liberal criteria for aura duration, 6 of 8 subjects who had been incorrectly classified as MwoA would instead be correctly classified as MwA by the POEM tool, increasing specificity for both MwA (94%) and MwoA (93%), and increasing sensitivity for MwA (90%). We have since modified the POEM questionnaire so that, if the subject indicates a typical aura duration of less than 5 or greater than 60 minutes, they are asked “Do these phenomena ever last 5-60 minutes?” An affirmative response is taken as consistent with a classification of MwA.

Other screening tools have been developed to identify patients with migraine in both clinical and research settings with varying test characteristics and degrees of accuracy (3–11); we summarize the properties of these in Table 3. Many of the tools, such as the ID Migraine (6), Migraine Screen Questionnaire (MS-Q) (9) and Migraine-4 (11), contain 3 to 5 questions to facilitate quick screening in the primary care setting. These clinical screening tools are generally sensitive but less specific (with the exception of the Migraine-4; (11). The Computerized Headache Assessment tool (CHAT) can identify several primary headache disorders including episodic and chronic migraine, but the sensitivities and specificities for the individual diagnoses have not been reported (22), and the tool itself is not available open source. The Structured Migraine Interview (SMI) is a 10-item questionnaire offered for both clinical and research purposes (12). While it has a high sensitivity, the SMI has a modest specificity of 0.58. The questions for SMI have been published, but the automated-algorithm does not appear to be open-source. Similar to the POEM, the deCODE Migraine Questionnaire (DMQ-3) is able to distinguish between MwoA and MwA (13), but the published link to the online questions for DMQ-3 is no longer available. Finally, the American Migraine Survey (AMS-II) is a 21-item questionnaire designed for epidemiological studies (23). The performance of the AMS-II for classification of migraine subtypes has not been examined, and publicly available analysis routines are not available.

**Table 3.**
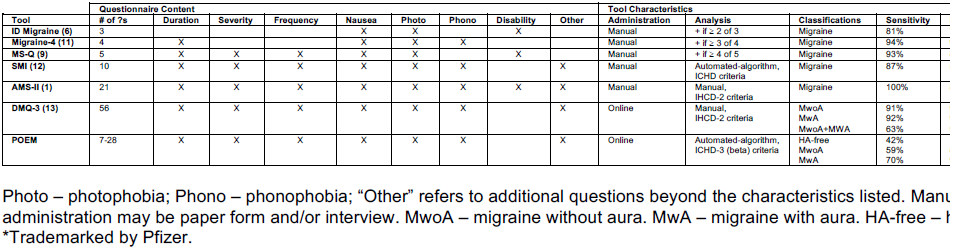
Comparison of migraine screening tools

These different instruments have been validated in various ways. Study populations have included clinical (6, 12) and non-clinical populations, with the latter composed of Pfizer employees (9), University Students (11), or the general US population (1). Sample sizes have ranged from 140 (9) to 942 (11) with a median of 200 subjects in each validation study. Five of the six studies used a clinical diagnosis by a neurologist as the gold standard, whereas the Migraine-4 (11) used the Standard-Diagnostic Interview for Headache-Revised (24).

In contrast to currently available tools, the POEM offers several advantages. First, the POEM not only identifies patients with migraine, but provides a sub-classification into migraine without aura, migraine with visual aura, and migraine with other aura. Second, the POEM is the only tool that specifically identifies headache-free subjects. Third, the POEM (like the Migraine-4) implements ICHD-3 (beta) criteria. Finally, the POEM is an open-source, automated, web-based tool, allowing for easy adoption by clinicians and researchers.

## Conclusions

The POEM is a new instrument to classify subjects into headache categories, including migraine with visual aura, migraine with other aura, migraine without aura, headache not otherwise specified, mild non-migrainous headache, and headache free. The migraine classifications meet ICHD-3 (beta) criteria (2). The tool is self-administered, avoiding the need for time- and cost-intensive interviews. The categorization is automated via open-source code. In our validation study, we find the POEM to have good specificity, and is thus suitable for the enrollment of subjects in migraine research trials.

## Institutional Review Board Approval

This study was approved by the University of Pennsylvania Institutional Review Board; all participants provided written informed consent.

## Article Highlights

- We have developed a self-administered, online survey and automated, open-source tool entitled the Penn Online Evaluation of Migraine (POEM) that provides six distinct headache classifications.
- To validate the POEM, 90 subjects completed the instrument and their classifications were compared to those given by a study neurologist during the ACHE-M study.
- While moderately sensitive, the POEM instrument is highly specific in classifying subjects into the categories of headache-free, migraine without aura, and migraine with aura, and is thus useful for enrolling subjects for research studies.

## Supplemental Tables/Figures

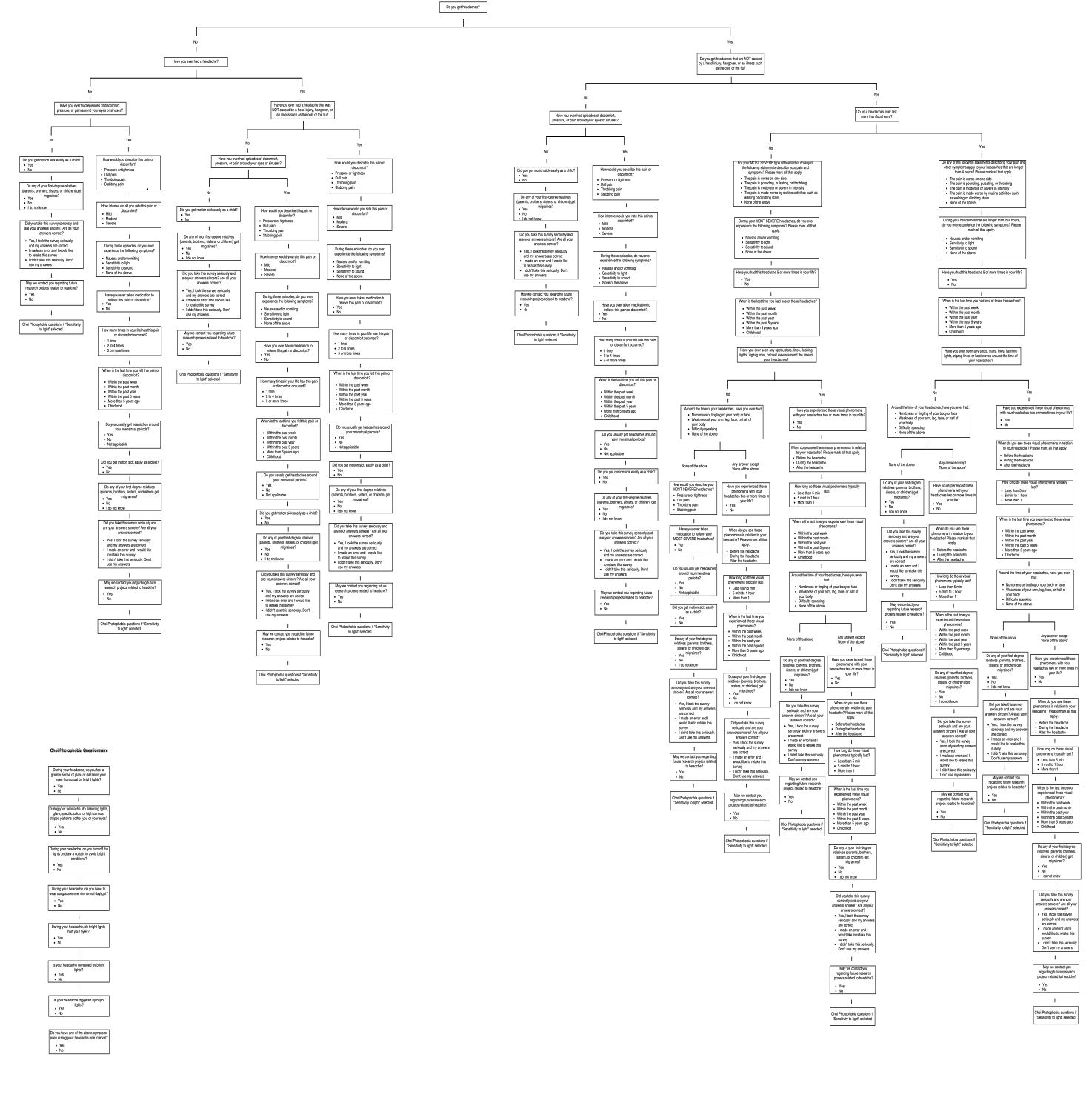

**Table S1.** Migraine without aura

| Question | Inclusion responses |
| --- | --- |
| 1. Do you get headaches? | Yes |
| 2. Do you get headaches that are NOT caused by a head injury, hangover, or an illness such as the cold or the flu? | Yes |
| 3. Do your headaches ever last more than 4 hours? | Yes |
| 4. Have you had this headache 5 or more times in your life? | Yes |
| 5. Do any of the following statements describing your pain and other symptoms apply to your headaches that are longer than 4 hours? Please mark all that apply. | – The pain is worse on one side<br>– The pain is pounding, pulsating, or throbbing<br>– The pain is moderate or severe in intensity<br>– The pain is made worse by routine activities such as walking or climbing stairs |
| 6. During your headaches that are longer than 4 hours, do you ever experience the following symptoms? Please mark all that apply. | – Nausea and/or vomiting<br>– Sensitivity to light<br>– Sensitivity to sound |

For migraine without aura, subjects need select at least 2 of 4 responses for question 5, and select 1 of 3 responses for question 6 in order to meet criteria.

**Table S2.** Migraine with visual aura

| Question | Inclusion responses |
| --- | --- |
| 1. Do you get headaches? | Yes |
| 2. Do you get headaches that are NOT caused by a head injury, hangover, or an illness such as the cold or the flu? | Yes |
| 3. Have you ever seen any spots, stars, lines, flashing lights, zigzag lines, or heat waves around the time of your headaches? | Yes |
| 4. Have you experienced these visual phenomena with your headaches two or more times in your life? | Yes |
| 5. How long do these visual phenomena typically last? | 5 min to 1 hour |
| 6. When do you see these visual phenomena in relation to your headache?<br>Please mark all that apply. | Before the headache |

For migraine with visual aura, subjects need to answer questions as stated.

**Table S3.** Migraine with other aura

| Question | Inclusion responses |
| --- | --- |
| 1. Do you get headaches? | Yes |
| 2. Do you get headaches that are NOT caused by a head injury, hangover, or an illness such as the cold or the flu? | Yes |
| 3. Around the time of your headaches, have you ever had: | – Numbness or tingling of your body or face<br>– Weakness of your arm, leg, face, or half of your body<br>– Difficulty speaking |
| 4. Have you experienced these phenomena with your headaches two or more times in your life? | Yes |
| 5. How long do these phenomena typically last? | – 5 min to 1 hour<br>– More than 1 hour |
| 6. When do you experience these phenomena in relation to your headache? Please mark all that apply. | Before the headache |

For migraine with other aura where individuals reported non-visual aura symptoms, subjects need to select at least 1 of 3 responses for question 3. Also, subject need to select either of the two responses listed for question 5, which is based on ICHD-3 (beta) criteria that state “each individual aura symptoms may last 5-60 min”; thus, subjects may have multiple aura symptoms that may in combination last longer than 60 min.

**Table S4.** Headache Free

|  |  |
| --- | --- |
| 1. Do you get headaches? | A. No; B. No; C. Yes |
| 2. Do you get headaches that are NOT caused by a head injury, hangover, or an illness such as the cold or the flu? | A. N/A; B. N/A; C. No |
| 3. Have you ever had a headache? | A. No; B. Yes; C. N/A |
| 4. Have you ever had a headache that was NOT caused by a head injury, hangover, or an illness such as the cold or the flu? | A. N/A; B. No; C. N/A |
| 5. Have you ever had episodes of discomfort, pressure, or pain around your eyes or sinuses? | A. No; B. No; C. No |

For headache free, subjects can meet criteria based on three potential pathways (A, B, or C) within the branching logic structure. Not applicable (N/A) questions were not in that specific pathway, thus, would not have been answered.

**Table S5.** Mild Non-Migrainous Headache

| Question | Exclusion responses |
| --- | --- |
| 1. Do your headaches ever last more than 4 hours? | Yes |
| 2. Have you ever seen any spots, stars, lines, flashing lights, zigzag lines, or heat waves around the time of your headaches? | Yes |
| 3. Around the time of your headaches, have you ever had: | – Numbness or tingling of your body or face<br>– Weakness of your arm, leg, face, or half of your body<br>– Difficulty speaking |
| 4. How would you describe this pain or discomfort? | – Throbbing pain<br>– Stabbing pain |
| 5. How intense would you rate this pain or discomfort | – Moderate<br>– Severe |
| 6. During these episodes, do you ever experience the following symptoms? Please mark all that apply. | – Nausea and/or vomiting<br>– Sensitivity to light<br>– Sensitivity to sound |
| 7. Do you usually get headaches around your menstrual periods? | – Yes |
| 8. For your MOST SEVERE type of headache, do any of the following statements describe your pain and symptoms? Please mark all that apply. | – The pain is pounding, pulsating, or throbbing<br>– The pain is moderate or severe in intensity<br>– The pain is made worse by routine activities such as walking or climbing stairs |
| 9. During your MOST SEVERE headaches, do you ever experience the following symptoms? Please mark all that apply | – Nausea and/or vomiting<br>– Sensitivity to light<br>– Sensitivity to sound |
| 10. How would you describe your MOST SEVERE headaches? | – Throbbing pain<br>– Stabbing pain |

To meet criteria for mild non-migrainous headache, subjects cannot select any of the responses listed above. Note that some questions and responses are slightly redundant due to the branching logic structure of the survey

## References

[1] Lipton RB, Bigal ME, Diamond M, Freitag F, Reed ML, Stewart WF. Migraine prevalence, disease burden, and the need for preventive therapy. Neurology. 2007;68(2007):343–9.

[2] Headache Classification Committee of the International Headache S. The International Classification of Headache Disorders, 3rd edition (beta version). Cephalalgia. 2013;33(2013):629–808.

[3] Michel P, Henry P, Letenneur L, Jogeix M, Corson A, Dartigues JF. Diagnostic screen for assessment of the IHS criteria for migraine by general practitioners. Cephalalgia. 1993;13 Suppl 12(1993):54–9.

[4] Gervil M, Ulrich V, Olesen J, Russell MB. Screening for migraine in the general population: validation of a simple questionnaire. Cephalalgia. 1998;18(1998):342–8.

[5] Pryse-Phillips W, Aube M, Gawel M, Nelson R, Purdy A, Wilson K. A headache diagnosis project. Headache. 2002;42(2002):728–37.

[6] Lipton RB, Dodick D, Sadovsky R, Kolodner K, Endicott J, Hettiarachchi J, et al. A self-administered screener for migraine in primary care: The ID Migraine validation study. Neurology. 2003;61(2003):375–82.

[7] Maizels M, Burchette R. Rapid and sensitive paradigm for screening patients with headache in primary care settings. Headache. 2003;43(2003):441–50.

[8] Cady RK, Borchert LD, Spalding W, Hart CC, Sheftell FD. Simple and efficient recognition of migraine with 3-question headache screen. Headache. 2004;44(2004):323–7.

[9] Lainez MJ, Dominguez M, Rejas J, Palacios G, Arriaza E, Garcia-Garcia M, Madrigal M. Development and validation of the Migraine Screen Questionnaire (MS-Q). Headache. 2005;45(2005):1328–38.

[10] Wang SJ, Fuh JL, Huang SY, Yang SS, Wu ZA, Hsu CH, et al. Diagnosis and development of screening items for migraine in neurological practice in Taiwan. Journal of the Formosan Medical Association = Taiwan yi zhi. 2008;107(2008):485–94.

[11] Walters AB, Smitherman TA. Development and Validation of a Four-Item Migraine Screening Algorithm Among a Nonclinical Sample: The Migraine-4. Headache. 2016;56(2016):86–94.

[12] Samaan Z, Macgregor EA, Andrew D, McGuffin P, Farmer A. Diagnosing migraine in research and clinical settings: the validation of the Structured Migraine Interview (SMI). BMC Neurol. 2010;10(2010):7.

[13] Kirchmann M, Seven E, Bjornsson A, Bjornssdottir G, Gulcher JR, Stefansson K, Olesen J. Validation of the deCODE Migraine Questionnaire (DMQ3) for use in genetic studies. European journal of neurology. 2006;13(2006):1239–44.

[14] Cucchiara B, Wolf RL, Nagae L, Zhang Q, Kasner S, Datta R, et al. Migraine with aura is associated with an incomplete circle of willis: results of a prospective observational study. PLoS One. 2013;8(2013):e71007.

[15] Choi JY, Kim YH, Oh K, Yu SW, Jung KY, Kim BJ. Cluster-like headache caused by posterior scleritis. Cephalalgia. 2009;29(2009):906–8.

[16] Headache CSotIHS. The International Classification of Headache Disorders: 2nd edition. Cephalalgia. 2004;24 Suppl 1(2004):9–160.

[17] Khil L, Straube A, Evers S, Berger K. Change in intraindividual ICHD-II headache diagnosis over time: a follow-up of the DMKG headache study. Cephalalgia. 2013;33(2013):25–33.

[18] Merikangas KR, Cui L, Richardson AK, Isler H, Khoromi S, Nakamura E, et al. Magnitude, impact, and stability of primary headache subtypes: 30 year prospective Swiss cohort study. Bmj. 2011;343(2011):d5076.

[19] Stang PE, Yanagihara PA, Swanson JW, Beard CM, O’Fallon WM, Guess HA, Melton LJ, 3rd. Incidence of migraine headache: a population-based study in Olmsted County, Minnesota. Neurology. 1992;42(1992):1657–62.

[20] Stewart WF, Linet MS, Celentano DD, Van Natta M, Ziegler D. Age- and sex-specific incidence rates of migraine with and without visual aura. American journal of epidemiology. 1991;134(1991):1111–20.

[21] Nachit-Ouinekh F, Chrysostome V, Henry P, Sourgen C, Dartigues JF, El Hasnaoui A. Variability of reported headache symptoms and diagnosis of migraine at 12 months. Cephalalgia. 2005;25(2005):117–23.

[22] Maizels M, Wolfe WJ. An expert system for headache diagnosis: the Computerized Headache Assessment tool (CHAT). Headache. 2008;48(2008):72–8.

[23] Lipton RB, Stewart WF, Diamond S, Diamond ML, Reed M. Prevalence and burden of migraine in the United States: data from the American Migraine Study II. Headache. 2001;41(2001):646–57.

[24] Andrew ME, Penzien DB, Rains JC, Knowlton GE, McAnulty RD. Development of a computer application for headache diagnosis: the Headache Diagnostic System. Int J Biomed Comput. 1992;31(1992):17–24.

